# Identification of replication fork-associated proteins in Drosophila embryos and cultured cells using iPOND coupled to quantitative mass spectrometry

**DOI:** 10.1101/2022.01.18.476773

**Authors:** Alexander Munden, Madison T. Wright, Dongsheng Han, Reyhaneh Tirgar, Lars Plate, Jared T. Nordman

## Abstract

Replication of the eukaryotic genome requires the formation of thousands of replication forks that must work in concert to accurately replicate the genetic and epigenetic information. Defining replication fork-associated proteins is a key step in understanding how genomes are replicated and repaired in the context of chromatin to maintain genome stability. To identify replication fork-associated proteins, we performed iPOND (Isolation of Proteins on Nascent DNA) coupled to quantitative mass spectrometry in Drosophila embryos and cultured cells. We identified 76 and 278 fork-associated proteins in post-MZT embryos and Drosophila cultured S2 cells, respectively. By performing a targeted screen of a subset of these proteins, we demonstrate that BRWD3, a targeting specificity factor for the DDB1/Cul4 ubiquitin ligase complex (CRL4), functions at replication forks to promote fork progression and maintain genome stability. Altogether, our work provides a valuable resource for those interested in the DNA replication, repair and chromatin assembly during development.

## Introduction

Each and every time a cell divides it must accurately replicate both its genetic and epigenetic information. Core replication factors are known to assemble at replication forks to replicate the genome (e.g. helicase, polymerases). There are, however, likely hundreds of proteins that function at or in close proximity to the replication fork to facilitate replication of difficult-to-replicate sequences, propagate epigenetic information and coordinate replication with other chromatin-related processes such as transcription^1–3^. Replication of the eukaryotic genome requires thousands of replication forks functioning simultaneously to complete replication in a timely manner. Errors generated at a single replication fork during genome duplication can result in mutations or genomic alterations with the potential to cause cell lethality or drive tumor formation^4^. Further complicating DNA replication is the need to allow regulatory flexibility to accommodate cell-type specific changes in cell division and cell cycle rates that occur during cell differentiation and development^5,6^. How replication fork composition and activity is remodeled in response to difficult-to-replicate regions of the genome and in response to changes in developmentally programed changes in S phase regulation has yet to be defined.

Drosophila provides an ideal system to understand how developmentally-programed changes in S phase impact replication fork composition. In contrast to the ∼8hr S phases associated with mitotic cell division in differentiated cells, S phases during early embryonic development are extremely rapid. During early embryogenesis and prior to the maternal-to-zygotic transition (MZT) S phases are 3-4 minutes in length. S phase gradually lengthens as development approaches the MZT and slows to ∼75 minutes at the MZT ^5,7^. While S phase length can drastically differ during development, the rate of replication fork progression is similar in these different contexts ^8^. The chromatin context that replication forks must navigate also change with development. In pre-MZT embryos, chromatin is devoid of heterochromatin and transcription is largely inactive. Around the time of the MZT, condensed heterochromatin is formed and zygotic transcription is activated^5,9^. In fact, the extension of S phase at the MZT is largely driven by the onset of late replication and the bulk of S phase is dedicated to replication of heterochromatic sequences^5,10^.

In recent years, several techniques have been established to isolate active replication forks to identify replication fork-associated proteins ^3,11,12^. One technique, isolation of proteins on nascent DNA (iPOND), has become widely employed due to its ease of use and the only technical requirement being a short pulse of the nucleotide analog 5-ethynyl-2'-deoxyuridine (EdU)^11^. For iPOND, cells are incubated with a brief pulse of EdU and proteins are crosslinked to nascent DNA. EdU can be biotinylated using click chemistry and newly synthesized DNA and associated proteins are purified using streptavidin beads^13^. A key to identifying proteins associated with active replication forks, rather than general chromatin-associated proteins, is a chase sample where a thymidine chase is introduced after the EdU pulse. Proteins enriched in pulse only samples relative to the chase samples are largely replication fork-associated proteins^2,11,14^. Another advantage of iPOND is that it can be coupled to quantitative mass spectrometry to identify replication fork-associated proteins in an unbiased manner^1,2,15^. While iPOND coupled to quantitative mass spectrometry has been used extensively in mammalian cultured cells, it has not been applied in Drosophila or in a developing organism.

To identify replication fork-associated proteins in Drosophila, and to determine if replication fork composition is influenced by development, we have performed iPOND in combination with tandem mass tag (TMT)-based quantitative mass spectrometry in Drosophila post-MZT embryos and cultured cells. Using an iPOND-TMT approach, together with a stringent statistical analysis, we identified 76 and 278 replication fork-associated proteins in post-MZT embryos and Drosophila cultured S2 cells, respectively. While we have confirmed many known replication fork components, we have identified many proteins that do not have known roles at the replication fork. By performing a targeted RNAi-based screen of select factors, we have identified the Cul4 E3 ubiquitin ligase specificity factor, BRWD3, as a replication fork-associated protein that affects replication fork progression.

## Results

### Establishing iPOND in the developing embryo

To define the landscape of replication fork-associated proteins during development, we turned to Drosophila due to its well-characterized S phase programs that are known to significantly change during development. To identify replication fork-associated proteins, we chose to use iPOND because it does not require protein tags or extensive multi-step purifications^11^. We chose post-MZT Drosophila embryos (3-5 hours after egg laying -AEL) for our embryonic sample. During this developmental time point, S phase is ∼ 75 minutes with the bulk of that time devoted to replication of heterochromatin (Fig. 1A)^5^.

**Figure 1.**
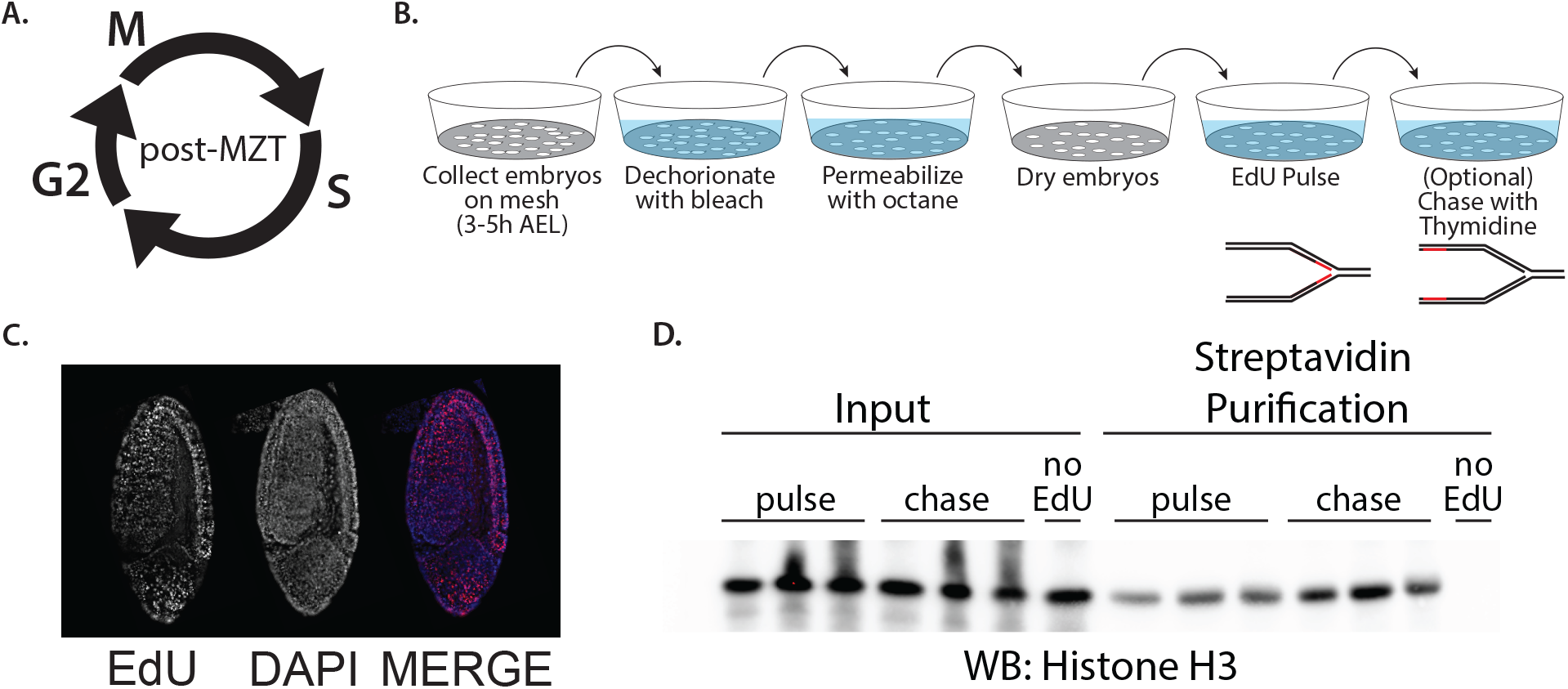
Establishing iPOND in the developing embryo. **(A)** A schematic of the rapid cell cycle in post-MZT embryos (3-5h AEL). **(B)** Experimental design for large-scale EdU labeling of embryos for iPOND. **(C)** Representative image of and EdU-labeled post-MZT embryo used for iPOND purifications. Embryo labeling was deemed successful if >50% of the embryos were uniformly labeled. **(D)** Western blot of three biological replicates for pulse and chase iPOND samples with a no EdU pulse control. Anti-H3 antibody is used as a marker of total chromatin recovery.

Nucleotide analogs and other small molecules are unable to enter embryos without permeabilization or direct injection^16^. To obtain sufficient EdU-labeled embryos for iPOND, we developed a large-scale permeabilization strategy. Starting with 3-5h embryos collected from a population cage, embryos were permeabilized and pulse-labeled with EdU for 10’ using custom collection baskets (see Experimental Procedures; Fig. 1B). Using this approach, we could routinely isolate 100-200mg of EdU-labeled embryos from a single collection basket. To determine if EdU-labeled 3-5h post-MZT embryos could be used for iPOND, we biotinylated embryos using Click chemistry as previously described ^13^ and the EdU-labeling efficiency was determined by staining embryos with fluorescently labeled streptavidin ^13^(Fig. 1C). Two key controls were also used; following the 10’ pulse of EdU, embryos were immediately transferred into medium containing thymidine for 30’ (chase sample). Second, age-matched embryos were mock treated and biotinylated exactly as the pulse and chase samples (no EdU control). To determine if iPOND could be used to isolate replication-dependent chromatin, we performed a single-step purification of biotinylated EdU-containing chromatin from our pulse, chase and no-EdU samples. We probed lysates for histone H3 as a mark of total chromatin^11^(Fig. 1D). We found that the recovery of chromatin was dependent on DNA replication. This indicates that iPOND can be applied to Drosophila embryos to isolate native chromatin.

### iPOND mass spectrometry identifies proteins associated with replication forks in Drosophila embryos

Now that we established iPOND as a technique to identify replication fork-associated proteins in Drosophila embryos, we wanted to identify the repertoire of proteins associated with replication forks in early embryos in an unbiased manner using quantitative mass spectrometry. Therefore, we coupled our iPOND purifications to tandem mass tag (TMT) labeling, which allows us to multiplex and quantify the relative abundance of peptides across multiple biological replicates in a single mass spectrometry experiment^17^. We optimized the amount of labeled embryos necessary for reproducible purification and mass spectrometry experiments. Ultimately, we found that 0.5g of EdU-labeled embryos routinely provided robust and reproducible mass spectrometry results. We collected EdU-pulsed embryos from four biological replicates of 3-5h embryos. EdU-pulsed embryos were either fixed immediately (pulse) or chased with thymidine for 30 minutes prior to fixation (chase). After EdU purification and verification via Western blot, peptides derived from pulse and chase samples were TMT labeled, separated using multidimensional protein identification technology (MudPIT) and quantified by mass spectrometry (Fig. 2A).

**Figure 2.**
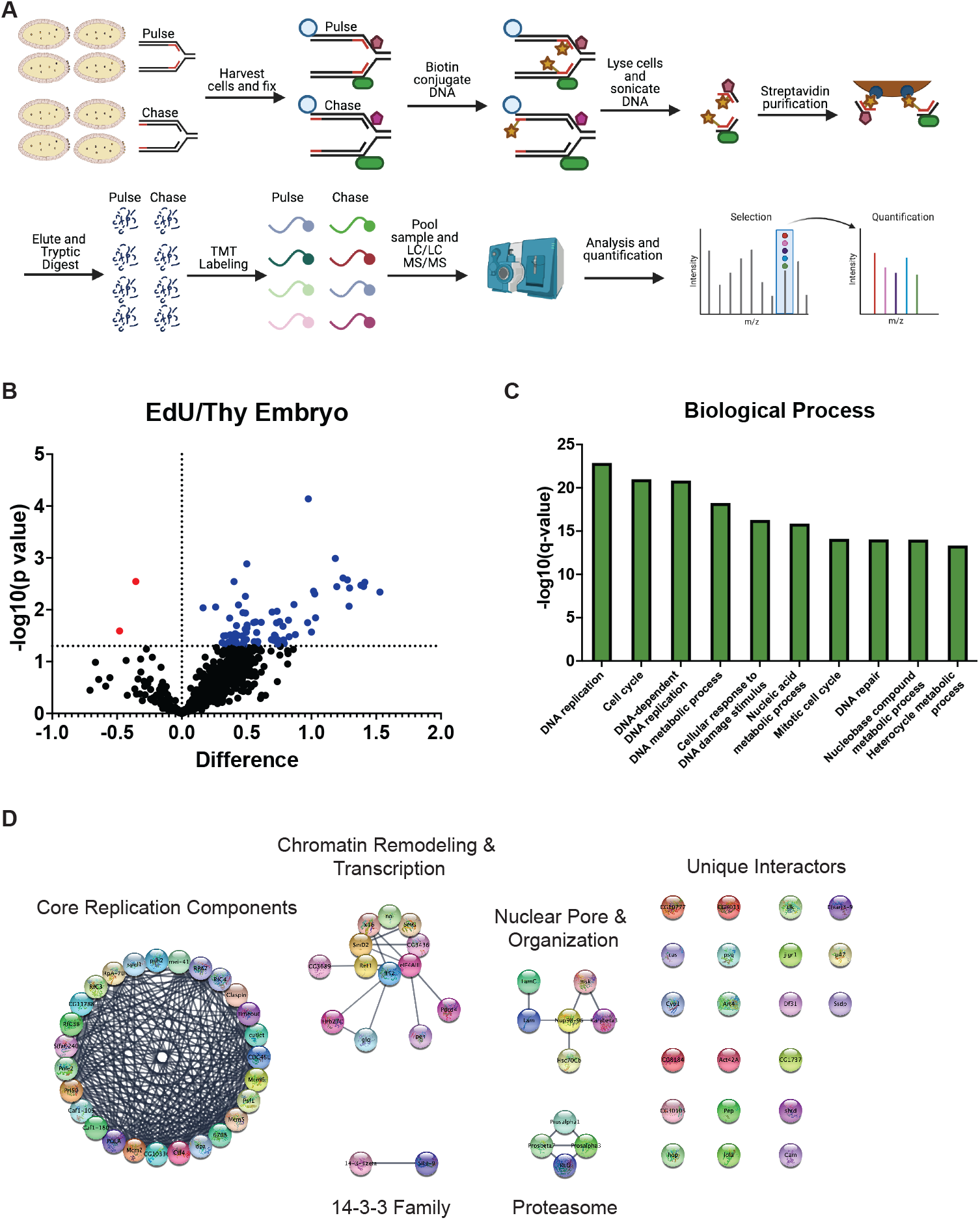
iPOND coupled to quantitative mass spectrometry in the post-MZT embryos. **(A)** A schematic of the labeling and mass spectrometry process for iPOND-TMT in Drosophila post-MZT embryos. **(B)** Volcano plot visualizing proteins identified as enriched or depleted in the pulse versus the chase embryo samples. Enrichment on the X-axis (log2[pulse]-log_2_[chase]) and -log_10_(p-value) on the Y-axis. **(C)** The top 10 enriched biological processes of the proteins enriched in the pulse sample as determined by Gene Ontology (GO) analysis. **(D)** Network map of the proteins enriched in the pulse sample, clustered into groups of known interactors using the STRING database with no additional interactors added.

To identify proteins that are enriched at replication fork s, we focused on proteins that were enriched in the EdU pulse samples relative to the thymidine chase controls. To ensure that the differences were not due to differences in purification and/or labeling efficiencies, we normalized proteins to histone H4 (see Methods). Using this analysis, we identified 76 proteins that were significantly enriched in either the EdU pulse or thymidine chase samples (Supp. Table 1). Most of the proteins enriched in our experiments were derived from the EdU pulse samples (Fig. 2B; 74 of 76). This is likely due to the relatively short thymidine chase time we adopted to allow us to focus on replication fork-associated proteins. Several lines of evidence indicate that our iPOND strategy is effective in isolating replication fork-associated proteins from Drosophila embryos. First, out of the top 25 enriched proteins in our data set, 18 are known replication factors (Supp. Table 2). Second, a Gene Ontology (GO) analysis of the 76 proteins enriched in the EdU pulse samples was highly enriched for DNA replication and DNA replication-associated processes (Fig. 2B). Therefore, we conclude that iPOND is an effective strategy to isolate replication fork-associated proteins during Drosophila embryogenesis.

We wanted to categorize the proteins in our data set in an unbiased manner to identify existing and potentially new protein networks centered around the replication machinery. To this end, we analyzed all enriched proteins using the STRING network in Cytoscape to identify proteins known to interact with one another. This analysis identified a network cluster of 27 known DNA replication and repair proteins (Fig. 2D). This cluster contains known helicase subunits, DNA polymerases, clamp loading factors and other factors (Fig. 2D). Additionally, it contains proteins involved in replication fork stability and response to DNA damage (Mei-41/ATR, Timeout/Timeless, CG10336/Tipin and Claspin)^18–22^. Unexpected clusters were also identified. For example, we identified a cluster of proteins involved in RNA processing and a cluster involved in nuclear organization and the nuclear pore. We also identified several unique replication fork-associated proteins that did not readily form interaction networks. Together, we conclude that iPOND can be used in Drosophila embryos to identify existing and potentially new replication fork-associated proteins.

### iPOND mass spectrometry identifies proteins associated with replication forks in Drosophila cultured cells

To extend the utility of iPOND in Drosophila, we performed iPOND in Drosophila S2 cultured cells. We previously performed iPOND in S2 cells, but did not couple iPOND to quantitative mass spectrometry^23^. First, we validated that iPOND functions in S2 cells (Supp. Fig. 1A). Next, we performed iPOND coupled to quantitative mass spectrometry with TMT labeling using ∼10^9^ cells/ biological replicate (Fig. 3A). Similar to our results in embryos, all of the enriched proteins were found in the pulse sample rather than the chase (Fig. 3B). This is likely due to the short chase time we used in these experiments. One difference we noted, however, is that we identified 278 replication fork-associated proteins in S2 cells (compared to 76 in embryos) (Supp. Table 3). While significantly higher than embryos, this protein number is similar to other iPOND and iPOND-like data sets in mammalian cells^1,3^.

**Figure 3.**
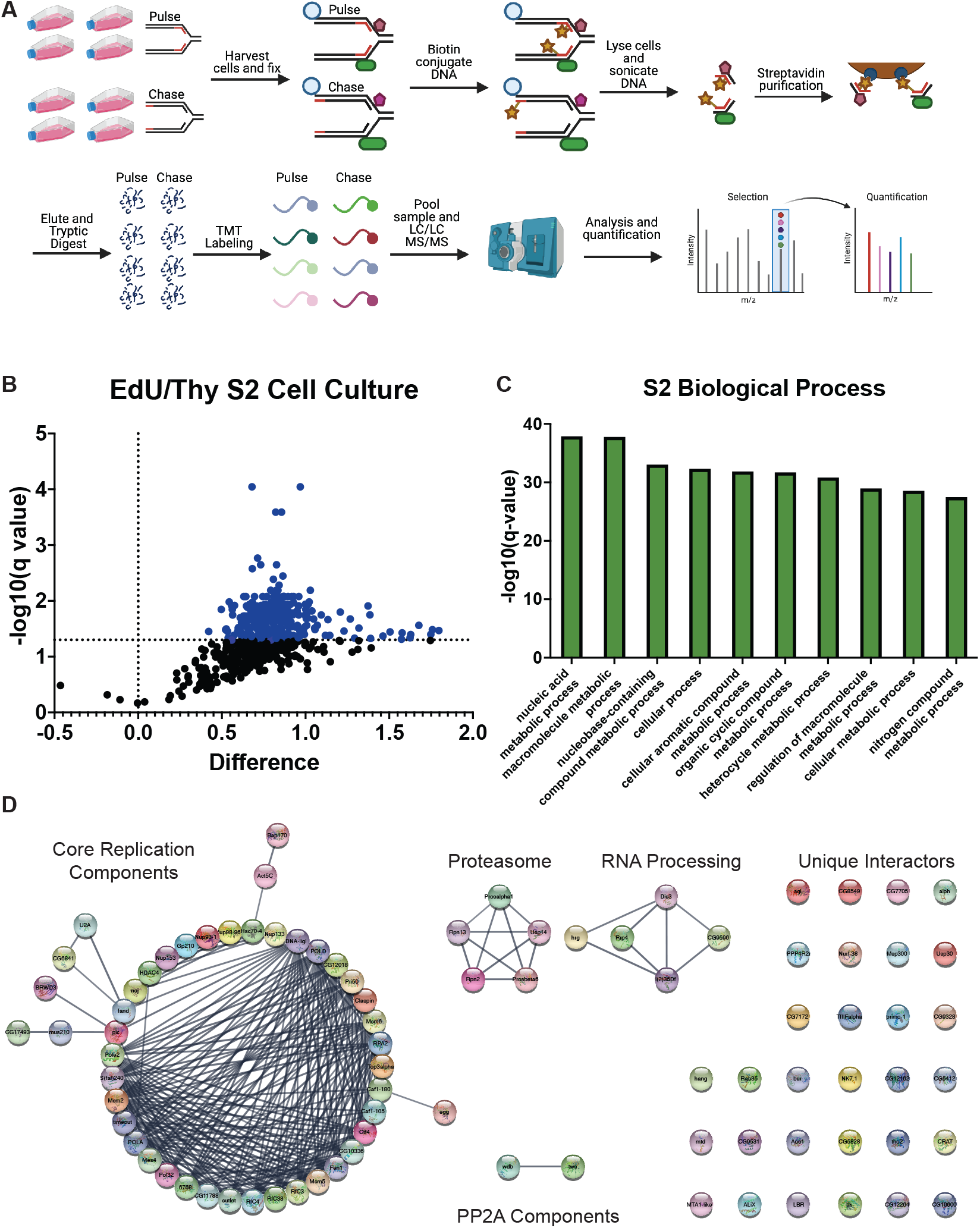
iPOND coupled to quantitative mass spectrometry in S2 cells. **(A)** A schematic of the labeling and mass spectrometry process for iPOND-TMT in Drosophila S2 cells. **(B)** Volcano plot visualizing those proteins identified as enriched or depleted in the pulse versus the chase cell culture samples. Enrichment on the X-axis (log2[pulse]-log_2_[chase]) on the X-axis and -log_10_(p-value) on the Y-axis. **(C)** The top 10 enriched biological processes of the proteins enriched in the pulse sample as determined by Gene Ontology (GO) analysis. **(D)** Network map of the proteins enriched in the pulse sample, clustered into groups of known interactors using the STRING database with no additional interactors added. For visualization, we included proteins with a corrected p value of <0.05 and a > 1.8-fold fold enrichment (99 total proteins). The full list of enriched interactors can be found in Supplemental Table 4.

Similar to our embryo data set, multiple lines of evidence indicate that our purifications successfully captured replication fork-associated proteins. Out of the top 25 enriched proteins in our data set, 22 are known replication factors (Supp. Table 4). Next, a Gene Ontology (GO) analysis of proteins enriched in the EdU pulse samples was highly enriched for DNA replication and DNA replication-associated processes (Fig. 3B). We attempted to generated an unbiased interaction network map using the STRING network in Cytoscape for the 278 enriched proteins, however, the networks were too dense to effectively visualize any meaningful interaction network hubs (Supp. Fig. 1B). For prioritization, we selected the proteins with an adjusted p value of < 0.05 and greater than 1.8-fold enrichment in the pulse relative to the chase. These stringent statistical cutoffs revealed 99 high-confidence proteins that were ultimately used to generate interaction network clusters, which consisted of known replication fork factors, further validating this data set and statistical analysis (Fig. 3D). Also, we identified networks containing proteosome components, RNA processing factors, protein phosphatase 4 complex (PP4) and a number of proteins with no recognized network connections (Fig. 3D)

### BRWD3 affects genome stability and replication fork progression

To determine if any of the replication fork-associated factors we identified affect genome stability, we used RNAi to deplete select factors and measured the global level of DNA damage. We chose to perform this targeted screen in S2 cells rather than post-MZT embryos due to the rapid and efficient depletion that can be obtained in S2 cells without the need to generate new reagents^24^. To quantify the global levels of DNA damage, we measured the level of phosphorylated H2Av (ɣ-H2Av), the Drosophila equivalent to mammalian ɣ-H2Ax, which is found at double strand breaks and stalled replication forks by immunofluorescence^25,26^. Of the 15 factors we chose, some but not all have known functions in DNA replication or DNA repair^18,27–35^. We validated knock down efficiency and the effect on cell proliferation for all factors (Supp. Fig 2A and 2B). As our negative control, we used a non-targeting RNA to *GFP* that is not present in S2 cells. As a positive control we targeted DNA polymerase alpha (*DNA pol?*), which is necessary for continual priming of the lagging strand (Fig 4a)^36^. Depletion of several factors resulted in increased H2Av phosphorylation (Fig. 4a). For example, depletion of *Cul4, RTEL, ELG1* and *BRWD3* all caused increased DNA damage consistent with mammalian studies^27–30^.

**Figure 4.**
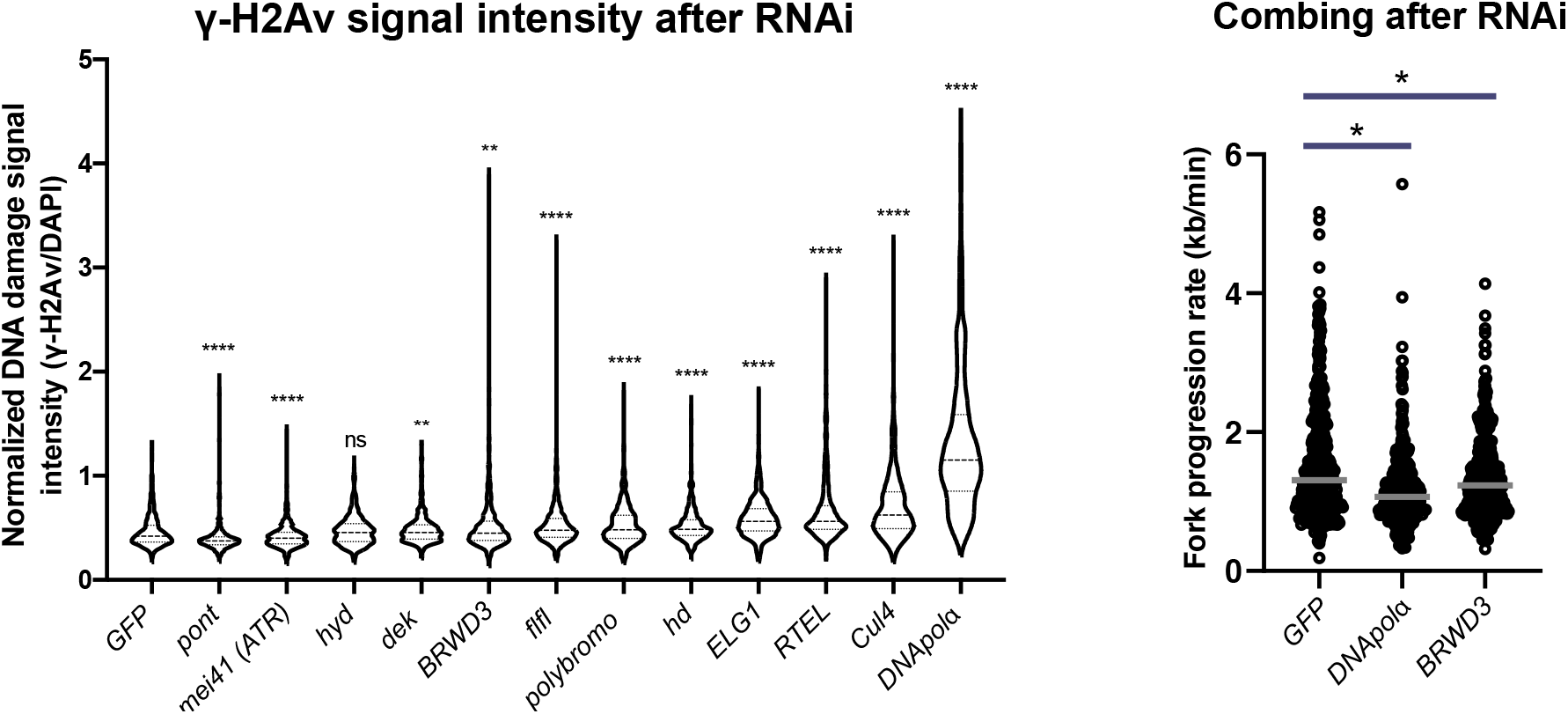
BRWD3 affects genome stability and replication fork progression. **(A)** RNAi-based depletion screen of candidate replication fork-associated proteins in S2 cells. Violin plots of the ɣ-H2Av intensity per nucleus normalized to total DNA content. Each distribution represents the signal intensities of 700 randomly selected cells from two biological replicates. **** p<0.0001 and ** p<0.01 using a Kruskal-Wallis one-way analysis of variance (B) Rate of fork progression in control and BRWD3 depleted S2 cells. 400 fibers from two biological replicates were pooled. Bars represent the median fork speed. * p<0.05 using a Kruskal-Wallis one-way analysis of variance followed by a Dunn’s multiple comparison post-test.

Interestingly, knockdown of *polybromo*, a component of the Brahma chromatin remodeling complex^37^, also caused an increase in DNA damage (Fig. 4a). Depletion of several factors caused a decrease in ɣ-H2Ax signal intensity, suggesting these factors contribute to DNA damage detection or signaling (Fig. 4a). Consistent with this hypothesis, depletion of *mei41* (the Drosophila ATR ortholog) decreased ɣ-H2Av intensity.

BRWD3 is a targeting specificity factor for the DDB1/Cul4 ubiquitin ligase complex (CRL4)^38^. In mammalian cells, one of the BRWD3 orthologs DCAF14/PHIP associates with replication forks upon DNA replication stress^29^. Depletion of DCAF14 results in a modest increase in DNA damage, which is exacerbated upon replication stress^29^. Depletion of BRWD3 in Drosophila S2 cells causes an increase in ɣ-H2Ax levels in unstressed cells (Fig. 4a). This suggests that BRWD3/DCAF14 has an evolutionarily conserved role at the replication fork to maintain genome stability. Given these observations, we wanted to determine if BRWD3 affects replication fork progression in unchallenged Drosophila cells. To this end, we developed a DNA combing protocol for Drosophila S2 cells. While we initially attempted to perform DNA combing with both IdU and CldU nucleoside analogs, we were unable to successfully perform combing with IdU (data not shown). This would have allowed us to measure fork rate, fork asymmetry and inter-origin distance. To solely measure the rate of fork progression, we performed DNA combing analysis with CldU as the sole nucleotide analog. As positive and negative controls, we used interfering RNAs against *DNA pol?* and *GFP*, respectively. Depletion of *BRWD3* caused a decreased rate of fork progression in untreated cells (Fig. 4b). Thus, we conclude that BRWD3 functions at the replication fork to promote replication fork progression and genome stability in Drosophila.

## Discussion

By developing a large-scale EdU labeling protocol in Drosophila embryos, we were able to perform iPOND in a developing organism. By coupling iPOND to quantitive mass spectrometry we identified 76 replication fork-associated proteins in Drosophila post-MZT embryos. Giving confidence to this method of identifying replication fork-associated proteins, 32 proteins we identified have known roles in DNA replication or repair. We note, however, that not all know replication fork-associated proteins were identified in our data set. Multiple reasons likely explain this observation. First, we used a stringent statistical cut off in our analysis to avoid false positives (see Methods). Second, either due to loss in purification or difficulty in mass spectrometry, some replication proteins are simply not detected by mass spectrometry, resulting in false negatives. Therefore, we suspect that 76 proteins are an underestimate of the total number of replication fork-associated proteins in post-MZT embryos.

While iPOND-TMT identified 76 proteins in post-MZT embryos, the same technique uncovered 278 proteins in Drosophila cultured cells. Although this number of proteins is higher than what we observed in embryos, it is similar to recent iPOND and iPOND-like experiments coupled to quantitative mass spectrometry performed in mammalian cells^1,3^. The difference in protein number between post-MZT embryos and cultured cells could be due to cell-type-specific factors in S2 cells or technical differences when performing iPOND in embryos vs. cultured cells. It should be noted, however, that the rate of replication fork progression is similar in Drosophila embryos and cultured cells^8^. Given that a single pulse of EdU is the only technical limitation with iPOND, it seems unlikely that differences in the amount of EdU labeling are responsible for the differences in protein number between the two developmental states. One complicating factor when trying to compare iPOND data sets for cell-type-specific factors is that lack of a protein in one sample could be due to a limitation in peptide detection in mass spectrometry. Therefore, we cannot use the lack of a protein in one developmental sample as direct evidence that a protein is cell-type specific. Nonetheless, our data reveal numerous replication fork-associated proteins in Drosophila embryos and cultured cells that can serve as a resource for anyone interested in replication fork composition and activity.

One factor that we identified as a replication fork-associated protein is BRWD3. Interestingly, one of the BRWD3 orthologs in mammalian cells also functions at the replication fork to maintain genome stability upon replication stress^29^. One key difference, however, is that in Drosophila BRWD3 functions at the replication fork in the absence of exogenous replication stress. Therefore, while BRWD3 and DCAF14 are both substrate specificity factors for CRL4, they likely function differently in mammalian cells and Drosophila. While it is tempting to speculate that CRL4^BRWD3^ targets a critical factor for ubiquitylation at the replication fork, further work will be necessary to test this hypothesis.

In summary, we have developed a protocol for the biochemical isolation of replication fork-associated proteins in Drosophila embryos and cultured cells. Our work suggests that replication fork composition can be modulated during development. Importantly, we have provided a resource of replication fork-associated factors in Drosophila for those interested in DNA replication, DNA repair and chromatin dynamics during replication.

### Experimental Procedures

#### EdU pulsing of embryos

Oregon R flies were expanded into population cages on grape juice plates supplemented with wet yeast. Cages were kept at 25°C in a humidified room and plates changed daily. Prior to embryo collections, flies were precleared for at least two hours. To acquire post-MZT embryos, flies were allowed to lay for two hours, and the plate was aged for three hours at 25°C to obtain 3-5 hours AEL embryos. Embryos were transferred to a container with a wire mesh bottom, washed in water and embryos were dechorionated in 50% bleach for two minutes. After washing, embryos were arranged in a monolayer on the mesh and bucket were dried with paper towels. Embryos were allowed to air dry 4-10 minutes, then submerged in octane for precisely 3.5 minutes with gentle shaking. Embryos were then air dried for one minute while shaking. Permeabilized embryos were pulsed with 10 μM EdU in EBR for 10 minutes. For chase samples, EdU-pulsed embryos were transferred to a new solution containing 20 μM of thymidine for an additional 30 minutes. After pulse/chasing, embryos were transferred to a scintillating flask in 10mL of heptane. 10mL of 4% PFA was added (2% final) and embryos were shaken vigorously at room temperature for 20 minutes. After fixation, the bottom layer of PFA was removed and an equal volume of methanol was added. Embryos were shaken by hand for one minute, settled and heptane was removed. Embryos were washed in methanol twice and transferred to PBS + 0.1% Triton X-100 and permeabilized overnight at 4°C. For each batch of embryos, a small fraction was taken and biotinylated and incubated with 568-Streptavidin to ensure that at least 50% of embryos were labeled. Successful collections were pooled to obtain 500 μL of embryos per biological replicate.

#### EdU pulsing of S2 cells

S2 cells were obtained directly from the DGRC. Cells were confirmed negative for mycoplasma contamination via PCR. Cells were grown in Schneider’s Drosophila Medium with 10% heat-inactivated FBS (Gemini Bio Products) and 100 U/mL of Penicillin/Streptomycin (Fisher Scientific) and kept at 25°C. Cells were pulsed as described in^13^. Briefly, three T225 flasks of 70% confluent cells were pulsed with 10 μM of EdU for 9 minutes. Cells were scraped and spun down for three minutes at 300xg. 10mL of 2% paraformaldehyde (PFA) was added to each flask and samples were fixed at room temperature on a nutator for 20 minutes. Paraformaldehyde was neutralized with glycine and cells were centrifuged for five minutes at 900xg at 4°C and resuspended in PBS with 0.1% Triton X-100 at 4°C until processing. For the chase sample, after centrifuging the cells were resuspended in cell media with 20 μM thymidine and incubated for 30 minutes in the cell culture incubator before fixation. Three T225 flasks were pooled for each replicate (∼7.5e^8^ cells per replicate).

#### iPOND

Embryos and S2 cells were biotinylated as described in^13^. Briefly, PBS, CuSO_4_, Biotin-Azide, and sodium ascorbate were mixed and added to labeled cells and embryos for 30 minutes. After biotinylation, cells or embryos were washed with PBS + 0.1% Triton X-100. A crude nuclear extract was generated by douncing embryos in Buffer 1 (15 mM HEPES pH 7.6, 10 mM KCl, 5 mM MgCl_2_, 0.1 mM EDTA, 0.5 mM EGTA, 350 mM sucrose)^39^ twelve times using a B-type homogenizer and centrifuged for 15 minutes at 8000xg. This pellet was resuspended in 1.2mL of LB3 (1 mM EDTA, 0.5 mM EGTA, 10 mM Tris pH 7.5, 100 mM NaCl, 0.1% Na-Deoxycholate, 0.5% N-Lauroyl sarcosine)^40^ with 2X protease inhibitors. Cells were resuspended in 1.2 mL LB3 lysis buffer with 2x protease inhibitors.

Samples were sonicated in a Bioruptor Plus (Diagenode) at high power, 10 cycles at 30” seconds on/30” seconds off. After a short break, samples were vortexed and this was repeated until 40 total cycles were achieved. 100 μL of Streptavidin C1 Dynabeads were extensively washed with LB3 and added to each sample. Samples were incubated at 4°C for two hours on a nutator. The unbound material was reserved to verify chromatin fragmentation. Beads were washed five times in LB3, with the 4^th^ wash containing 500 mM NaCl. To elute, samples were incubated at 65°C overnight on thermoblock in 1:1 combination of LB3:SB (20% glycerol, 20% SDS, 120 mM Tris pH 6.8). The next day, the eluate was removed from the beads and added to 2x Laemmli buffer with DTT and boiled for 10 minutes. This lysate was used for western blot and mass spectrometry experiments.

#### Western blotting

Lysates from iPOND samples were loaded onto a 4-15% Mini-Protean Stain-free protein gel (BioRad). After running the gel, samples were transferred onto 0.2 μM PVDF using the Transblot Turbo system (BioRad). Membranes were blocked in 5% milk, and incubated with the appropriate antibody for 1 hour at room temperature. Histone H3 (abcam 21054, 1:3000) was used to verify the success of iPOND. After washing in TBS + 0.1% Tween-20 (TBST), secondary antibodies (Jackson Labs) conjugated with HRP were added at 1:10,000 (mouse) or 1:20,000 (rabbit). After 30 minutes at room temperature, membranes were washed with TBST, incubated with Clarity ECL for 5 minutes (Bio-Rad) and visualized using a Bio-Rad ChemiDoc MP Imaging System.

#### TMT Labeling

After verifying iPOND was successful by Western blot (5% of total material), the remaining purified material was precipitated using methanol and chloroform and washed with methanol to remove excess detergent. Protein was resuspended in 5 μL fresh 1% Rapigest. 32.5 μL of mass spectrometry grade water with HEPES (pH 8.0 at a final concentration of 100mM). Disulfide bonds were reduced with freshly made 5mM TCEP and incubated for 30 minutes at room temperature. Fresh Iodoacetamide was added at a final concentration of 10mM to acetylate free sulfhydryl bonds. Protein was digested overnight with 0.5 μg trypsin at 37°C with shaking and covered from light. The next day, samples were labeled using a TMT10plex kit (Thermo Scientific catalog #90110). TMT labels were resuspended in acetonitrile and each sample was incubated with the appropriate amount of TMT reagent for 1 hour at room temperature. Excess label was neutralized with 0.4% final concentration of ammonium bicarbonate for 1 hour. Samples were mixed and acidified with formic acid to a pH 2. The mixed sample was reduced to 1/6 of the original volume using a SpeedVac, and brought back up to original volume with Buffer A (5% acetonitrile, 0.1% formic acid). Rapigest was cleaved by incubating for one hour at 42°C. The samples were centrifuged at 14,000 rpm for 30 minutes and the supernatant was transferred to a fresh tube and stored at -80°C until mass spectrometry analysis.

#### Liquid Chromatography – Tandem Mass Spectrometry

MudPIT microcolumns were prepared as previously described^41^. Peptide samples were directly loaded onto the columns using a high-pressure chamber. Samples were then desalted for 30 minutes with buffer A (97% water, 2.9% acetonitrile, 0.1% formic acid v/v/v). LC-MS/MS analysis was performed using a Q-Exactive HF (Thermo Fisher) or Exploris480 (Thermo Fisher) mass spectrometer equipped with an Ultimate3000 RSLCnano system (Thermo Fisher). Embryo samples were analyzed on the Exploris480 while the S2 cell culture were analyzed on the Q-Exactive HF. MudPIT experiments were performed with 10μL sequential injections of 0, 10, 30, 60, and 100% buffer C (500mM ammonium acetate in buffer A), followed by a final injection of 90% buffer C with 10% buffer B (99.9% acetonitrile, 0.1% formic acid v/v) and each step followed by a 130 minute gradient from 5% to 80% B with a flow rate of 300nL/minute when using the Q-Exactive HF and 500nL/minute when using the Exploris480 on a 20cm fused silica microcapillary column (ID 100 um) ending with a laser-pulled tip filled with Aqua C18, 3μm, 100 Å resin (Phenomenex). Electrospray ionization (ESI) was performed directly from the analytical column by applying a voltage of 2.0kV when using the Q-Exactive HF and 2.2kV when using the Exploris480 with an inlet capillary temperature of 275°C. Using the Q-Exactive HF, data-dependent acquisition of mass spectra was carried out by performing a full scan from 300-1800 m/z with a resolution of 60,000. The top 15 peaks for each full scan were fragmented by HCD using normalized collision energy of 38, 0.7 m/z isolation window, 120 ms maximum injection time, at a resolution of 45,000 scanned from 100 to 1800 m/z and dynamic exclusion set to 60s. Using the Exploris480, data-dependent acquisition of mass spectra was carried out by performing a full scan from 400-1600m/z at a resolution of 120,000. Top-speed data acquisition was used for acquiring MS/MS spectra using a cycle time of 3 seconds, with a normalized collision energy of 36, 0.4m/z isolation window, 120ms maximum injection time, at a resolution of 45,000 with the first m/z starting at 110. Peptide identification and TMT-based protein quantification was carried out using Proteome Discoverer 2.4. MS/MS spectra were extracted from Thermo Xcalibur .raw file format and searched using SEQUEST against a Uniprot *Drosophila melanogaster* proteome database (downloaded February 6^th^, 2019 and containing 21114 entries). The database was curated to remove redundant protein and splice-isoforms, and supplemented with common biological MS contaminants. Searches were carried out using a decoy database of reversed peptide sequences and the following parameters: 10ppm peptide precursor tolerance, 0.02 Da fragment mass tolerance, minimum peptide length of 6 amino acids, trypsin cleavage with a maximum of two missed cleavages, dynamic methionine modification of 15.995 Da (oxidation), static cysteine modification of 57.0215 Da (carbamidomethylation), and static N-terminal and lysine modifications of 229.1629 Da.

#### iPOND-TMT data analysis

To determine enrichment or depletion of the proteins, the TMT intensities for each protein was log_2_ transformed and samples were normalized based on median TMT intensity per channel. Log2-transformed, median normalized TMT intensities were further normalized to the level of Histone H4, as the resulting incorporation of this histone should be identical between each sample. Enrichment values were calculated based on this normalized data. Cellular localization data was determined for each protein using the Gene Ontology Cellular Compartment (FlyBase v2021_05). Proteins that lacked any nuclear or chromatin compartmental data were removed from the datasets. To determine if a protein was significantly enriched or depleted in the pulse or chase embryo samples, an unpaired t-test was performed for each protein. Our uncorrected p values were validated because our positive controls (known replication proteins) were identified. For S2 cell data, an unpaired t-test was performed for each protein with a Benjamini, Krieger and Yekutieli multiple test correction and false discovery rate of 5%^42^.

For the pathway enrichment analysis of enriched proteins, PANTHER Gene Ontology was used ^43–45^. Enriched proteins were inputted and the default background for *Drosophila melanogaster* was selected. The biological process pathway was used, and the results were exported to Excel and the top 10 pathways were chosen by q-value, and visualized in Graphpad Prism

For network clustering, all of the proteins enriched in the embryo and S2 pulse were loaded as separate networks in Cytoscape v3.9.0^46^. The resulting interactions were visualized using the STRING network with the stringApp, using the *Drosophila melanogaster* setting with 0 additional interactors and a confidence score cutoff of 0.8^47,48^.

#### RNAi and immunofluorescence in S2 cells

RNAi in S2 cells was performed as described^49^. Briefly, dsRNA against each candidate RNA was designed to be 200-500 bp. The dsRNAs were synthesized using the Invitrogen MEGAscript T7 Transcription Kit (Ambion). For each sample, 1.5 million S2 cells were seeded in 1 ml non-serum medium in a 6-well plate and 30 μg of dsRNA was added. After 45 minutes incubation at room temperature, 2ml of serum-containing medium was added and cells were incubated for an additional five days. Reverse transcription and quantitative PCR (RT-qPCR) was performed to determine the knock down efficiency. For immunofluorescence, RNAi-treated cells were attached to Concanvan A-coated slides for 15 minutes, fixed for 15 minutes in 4% paraformaldehyde and permeabilized for 15 minutes in PBS supplemented with 0.3% Triton-X-100 (PBT). Cells were then blocked for 60 minutes in blocking buffer, containing 1% BSA and 0.2% goat serum in 0.1% PBT. After blocking, cells were incubated with rabbit anti-γ-H2Av (1:500, Rockland, # 600-401-914) antibody overnight at 4°C in blocking buffer. After washing with PBS, cells were incubated with goat anti-rabbit IgG secondary antibody (1:500, Life Technologies, # A11011) in blocking buffer for one hour at room temperature and stained with DAPI (0.1 μg/mL) in PBT for 10 minutes and mounted in Vectasheild (Vector Labs). All images were obtained using Nikon Ti-E inverted microscope with a Zyla sCMOS digital camera with a 20X oil objective. For each biological replicate, all samples were captured at the same magnification and same exposure time. For quantitative analysis of γ-H2Av levels, regions of interest (ROIs) were defined based on the DAPI signal. The mean signal intensity of ɣ-H2Av was extracted for each ROI. The signal was normalized to the DAPI signal intensity to account for differences in the total amount of DNA. 350 randomly selected cells were used for each biological replicate. Two biological replicates were used for the data analysis. Kruskal-Wallis one-way analysis of variance was performed in GraphPad Prism for statistical significance.

#### DNA molecular combing

Drosophila S2 cells were pulsed with 20uM of CldU nucleoside (Sigma-Aldrich, C6891) for 20 minutes. Cells were washed with PBS then ∼1.5-3.0 million cells were embedded in agarose plugs. The assay was performed as described in Genomic Vision’s manufacturer instructions. The stretched and denatured DNA was stained with a CldU-specific antibody (Abcam Cat#ab6326) for 1 hour, washed in PBS, then probed with a secondary antibody (Thermo, A11007) for 30min. Coverslips were washed with PBS then mounted. Stained coverslips were imaged using a Nikon Ti-E inverted microscope with a Zyla sCMOS digital camera with a 40X oil objective. For each sample, 200 DNA fiber lengths were measured manually using Nikon NIS-Elements AR v4.40. Investigator was blinded to sample identity. Two biological replicates were performed per sample. The length of a given fiber is directly proportional to the rate of replication fork progression. Therefore, fiber lengths were converted to fork progression rates given the 20-minute pulse time (Fiber length *20min / 2kb•min^-1^). Kruskal-Wallis one-way analysis of variance followed by Dunn’s multiple comparisons post-test was performed in GraphPad Prism for statistical significance.

## Acknowledgments

We thank Martina Brienza-Ramos for assistance with EdU labeling of embryos used in this study. We thank David Cortez and Kavi Mehta for advice and guidance with DNA combing. We thank Sarah Wessel and members of the Nordman lab for providing critical feedback on the manuscript. This work was supported by National Institutes of Health (NIH) General Medical Sciences awards R35GM133552 to L.P. and R35GM128650 to J.T.N.. M.T.W was supported by the Vanderbilt Chemistry-Biology Interface Training Program (T32GM065086) and the National Science Foundation Graduate Research Fellowship Program.

## Author Contributions

A.M. and J.T.N planned and designed the research; A.M., R.T, D.S performed experiments; A.M., M.W, D.H., and R.T. analyzed data with supervision from L.P and J.T.N; A.M. and J.T.N. wrote the manuscript with input from all of the authors. A.M., M.W., D.H., R.T., L.P. and J.T.N. edited the manuscript.

## Additional Information

### Supplementary information

Two supplemental figures and four supplemental tables

### Competing interest

The authors declare no competing interests

## Notes

### Competing Interest Statement

The authors have declared no competing interest.

## References

1. Wessel, S. R., Mohni, K. N., Luzwick, J. W., Dungrawala, H. & Cortez, D. Functional Analysis of the Replication Fork Proteome Identifies BET Proteins as PCNA Regulators. Cell Reports 28, 3497-3509.e4 (2019).

2. Dungrawala, H. et al. The Replication Checkpoint Prevents Two Types of Fork Collapse without Regulating Replisome Stability. Mol Cell 59, 998–1010 (2015).

3. Alabert, C. et al. Nascent chromatin capture proteomics determines chromatin dynamics during DNA replication and identifies unknown fork components. Nat Cell Biol 16, 281–291 (2014).

4. Tomasetti, C., Li, L. & Vogelstein, B. Stem cell divisions, somatic mutations, cancer etiology, and cancer prevention. Science 355, 1330–1334 (2017).

5. Shermoen, A. W., McCleland, M. L. & O’Farrell, P.H. Developmental Control of Late Replication and S Phase Length. Curr Biol 20, 2067–2077 (2010).

6. Matson, J. P. et al. Rapid DNA replication origin licensing protects stem cell pluripotency. eLife 6, e30473 (2017).

7. Yuan, K., Seller, C. A., Shermoen, A. W. & O’Farrell, P. H. Timing the Drosophila Mid-Blastula Transition: A Cell Cycle-Centered View. Trends Genet 32, 496–507 (2016).

8. Blumenthal, A. B., Kriegstein, H. J. & Hogness, D. S. The Units of DNA Replication in Drosophila melanogaster Chromosomes. Cold Spring Harb Sym 38, 205–223 (1974).

9. Tadros, W. & Lipshitz, H. D. The maternal-to-zygotic transition: a play in two acts. Development 136, 3033–3042 (2009).

10. Seller, C. A. & O’Farrell, P. H. Rif1 prolongs the embryonic S phase at the Drosophila mid-blastula transition. PLoS Biol 16, e2005687 (2018).

11. Sirbu, B. M. et al. Analysis of protein dynamics at active, stalled, and collapsed replication forks. Genes & Dev 25, 1320–1327 (2011).

12. Gambus, A. et al. GINS maintains association of Cdc45 with MCM in replisome progression complexes at eukaryotic DNA replication forks. Nat Cell Biol 8, 358–366 (2006).

13. Dungrawala, H. & Cortez, D. The Nucleus. Methods Mol Biology 1228, 123–131 (2014).

14. Sirbu, B. M. et al. Identification of Proteins at Active, Stalled, and Collapsed Replication Forks Using Isolation of Proteins on Nascent DNA (iPOND) Coupled with Mass Spectrometry. J Biol Chem 288, 31458–31467 (2013).

15. Cortez, D., Proteomic Analyses of the Eukaryotic Replication Machinery. Methods Enzymol 591, 33–53 (2017).

16. Limbourg, B. & Zalokar, M. Permeabilization of Drosophila eggs. Dev Biol 35, 382–387 (1973).

17. McAlister, G. C. et al. Increasing the Multiplexing Capacity of TMTs Using Reporter Ion Isotopologues with Isobaric Masses. Anal Chem 84, 7469–7478 (2012).

18. Sibon, O. C. M., Laurençon, A., Hawley, R. S. & Theurkauf, W. E. The Drosophila ATM homologue Mei-41 has an essential checkpoint function at the midblastula transition. Curr Biol 9, 302–312 (1999).

19. Benna, C. et al. Drosophila timeless2 Is Required for Chromosome Stability and Circadian Photoreception. Curr Biol 20, 346–352 (2010).

20. Lee, E.-M. et al. Drosophila Claspin is required for the G2 arrest that is induced by DNA replication stress but not by DNA double-strand breaks. DNA Repair 11, 741–752 (2012).

21. Gotter, A. L., Suppa, C. & Emanuel, B. S. Mammalian TIMELESS and Tipin are Evolutionarily Conserved Replication Fork-associated Factors. J Mol Biol 366, 36–52 (2007).

22. Saldivar, J. C., Cortez, D. & Cimprich, K. A. The essential kinase ATR: ensuring faithful duplication of a challenging genome. Nat Rev Mol Cell Bio 18, 622–636 (2017).

23. Munden, A. et al. Rif1 inhibits replication fork progression and controls DNA copy number in Drosophila. eLife 7, e39140 (2018).

24. Echeverri, C. J. & Perrimon, N. High-throughput RNAi screening in cultured cells: a user’s guide. Nat Rev Genet 7, 373–384 (2006).

25. Madigan, J. P., Chotkowski, H. L. & Glaser, R. L. DNA double-strand break-induced phosphorylation of Drosophila histone variant H2Av helps prevent radiation-induced apoptosis. Nucleic Acids Res 30, 3698–3705 (2002).

26. Mah, L.-J., El-Osta, A. & Karagiannis, T. C. γH2AX: a sensitive molecular marker of DNA damage and repair. Leukemia 24, 679–686 (2010).

27. Jin, J., Arias, E. E., Chen, J., Harper, J. W. & Walter, J. C. A Family of Diverse Cul4-Ddb1-Interacting Proteins Includes Cdt2, which Is Required for S Phase Destruction of the Replication Factor Cdt1. Mol Cell 23, 709–721 (2006).

28. Vannier, J.-B. et al. RTEL1 Is a Replisome-Associated Helicase That Promotes Telomere and Genome-Wide Replication. Science 342, 239–242 (2013).

29. Townsend, A., Lora, G., Engel, J., Tirado-Class, N. & Dungrawala, H. DCAF14 promotes stalled fork stability to maintain genome integrity. Cell Reports 34, 108669 (2021).

30. Bell, D. W. et al. Predisposition to Cancer Caused by Genetic and Functional Defects of Mammalian Atad5. PLoS Genet 7, e1002245 (2011).

31. Bandura, J. L. et al. humpty dumpty Is Required for Developmental DNA Amplification and Cell Proliferation in Drosophila. Curr Biol 15, 755–759 (2005).

32. Mansfield, E., Hersperger, E., Biggs, J. & Shearn, A. Genetic and molecular analysis of hyperplastic discs, a gene whose product is required for regulation of cell proliferation in Drosophila melanogaster imaginal discs and germ cells. Dev Biol 165, 507–26 (1994).

33. Kappes, F. et al. The DEK oncoprotein is a Su(var) that is essential to heterochromatin integrity. Genes & Dev 25, 673–678 (2011).

34. Sousa-Nunes, R., Chia, W. & Somers, W. G. Protein Phosphatase 4 mediates localization of the Miranda complex during Drosophila neuroblast asymmetric divisions. Genes & Dev 23, 359– 372 (2009).

35. Klymenko, T. et al. A Polycomb group protein complex with sequence-specific DNA-binding and selective methyl-lysine-binding activities. Genes & Dev 20, 1110–1122 (2006).

36. Muzi-Falconi, M., Giannattasio, M., Foiani, M. & Plevani, P. The DNA Polymerase alpha-Primase Complex: Multiple Functions and Interactions. Sci World J 3, 21–33 (2003).

37. Thompson, M. Polybromo-1: The chromatin targeting subunit of the PBAF complex. Biochimie 91, 309–319 (2009).

38. Jackson, S. & Xiong, Y. CRL4s: the CUL4-RING E3 ubiquitin ligases. Trends in Biochem Sci 34, 562–570 (2009).

39. Shao, Z. et al. Stabilization of Chromatin Structure by PRC1, a Polycomb Complex. Cell 98, 37–46 (1999).

40. MacAlpine, H. K., Gordân, R., Powell, S. K., Hartemink, A. J. & MacAlpine, D. M. Drosophila ORC localizes to open chromatin and marks sites of cohesin complex loading. Genome Res 20, 201–211 (2010).

41. Fonslow, B. R. et al. Single-Step Inline Hydroxyapatite Enrichment Facilitates Identification and Quantitation of Phosphopeptides from Mass-Limited Proteomes with MudPIT. J Proteome Res 11, 2697–2709 (2012).

42. Storey, J. D. & Tibshirani, R. Statistical significance for genomewide studies. PNAS 100, 9440–9445 (2003).

43. Ashburner, M. et al. Gene Ontology: tool for the unification of biology. Nat Genet 25, 25–29 (2000).

44. Carbon, S. et al. The Gene Ontology resource: enriching a GOld mine. Nucleic Acids Res 49, D325–D334 (2020).

45. Mi, H., Muruganujan, A., Ebert, D., Huang, X. & Thomas, P. D. PANTHER version 14: more genomes, a new PANTHER GO-slim and improvements in enrichment analysis tools. Nucleic Acids Res 47, D419–D426 (2019).

46. Shannon, P. et al. Cytoscape: A Software Environment for Integrated Models of Biomolecular Interaction Networks. Genome Res 13, 2498–2504 (2003).

47. Szklarczyk, D. et al. The STRING database in 2021: customizable protein-protein networks, and functional characterization of user-uploaded gene/measurement sets. Nucleic Acids Res 49, D605–D612 (2021).

48. Doncheva, N. T., Morris, J. H., Gorodkin, J. & Jensen, L. J. Cytoscape StringApp: Network Analysis and Visualization of Proteomics Data. J Proteome Res 18, 623–632 (2019).

49. Rogers, S. L. & Rogers, G. C. Culture of Drosophila S2 cells and their use for RNAi-mediated loss-of-function studies and immunofluorescence microscopy. Nat Protoc 3, 606–611 (2008).

