## Supplemental Figures for "Identification of replication fork-associated proteins in Drosophila embryos and cultured cells using iPOND coupled to quantitative mass spectrometry"

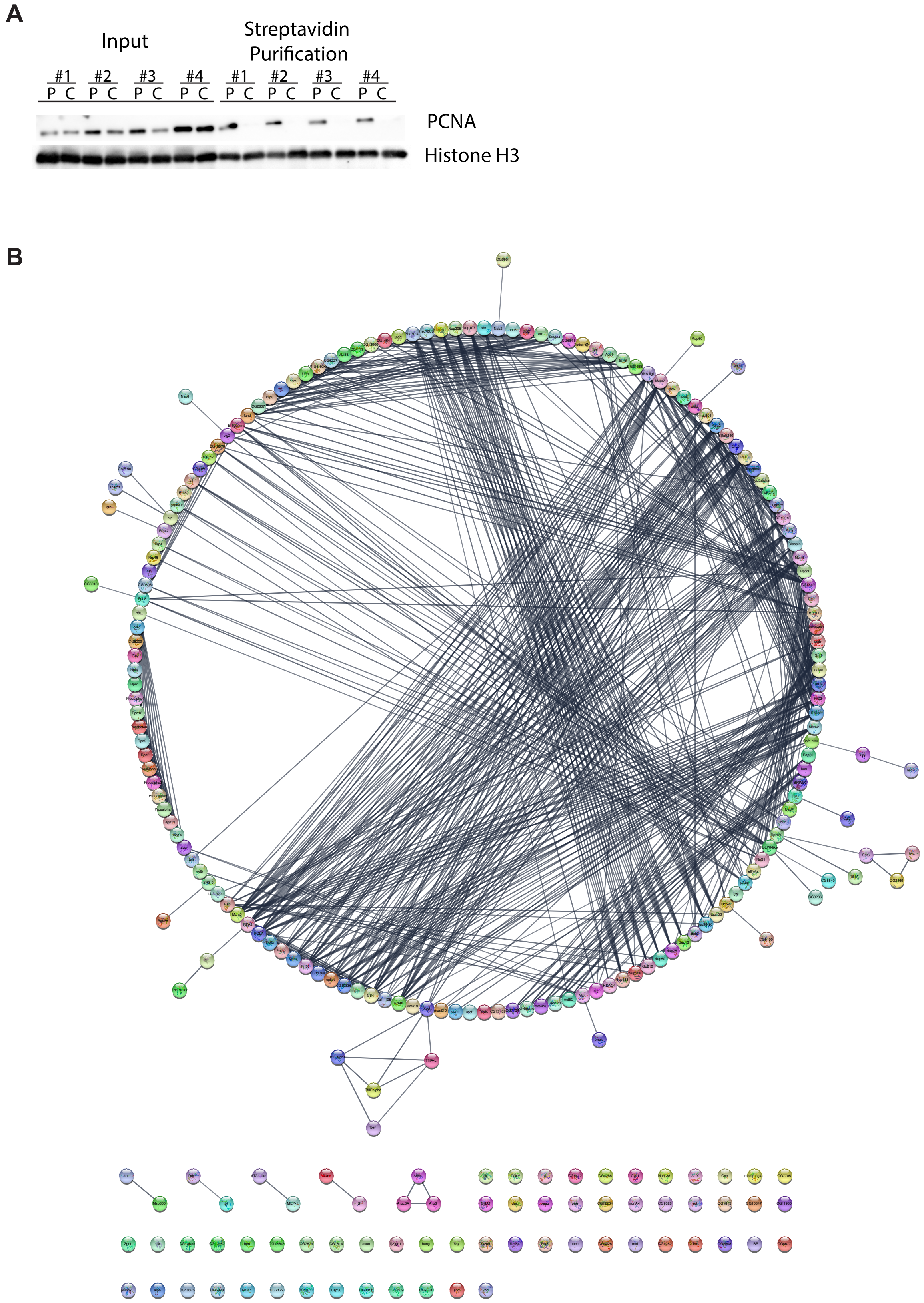
A
Input
Streptavidin Purification
#1 #2 #3 #4 #1 #2 #3 #4
P C P C P C P C P C P C P C P C
PCNA
Histone H3
B

**Supplemental Figure 1.** Expanded iPOND confirmation and findings in S2 cultured cells. (**A**) Western blot confirmation of iPOND in cell culture cells. The first 8 lanes are from input, and the last half from the streptavidin purification. PCNA is a marker for active replication forks and is only enriched in the pulse sample purifications as expected, while histone H3 is enriched in both the pulse and chase. P = Pulse, C = Chase, and #1-4 represent the replicate numbers. (**B**) Total network map of all proteins enriched in the pulse for cell culture cells.

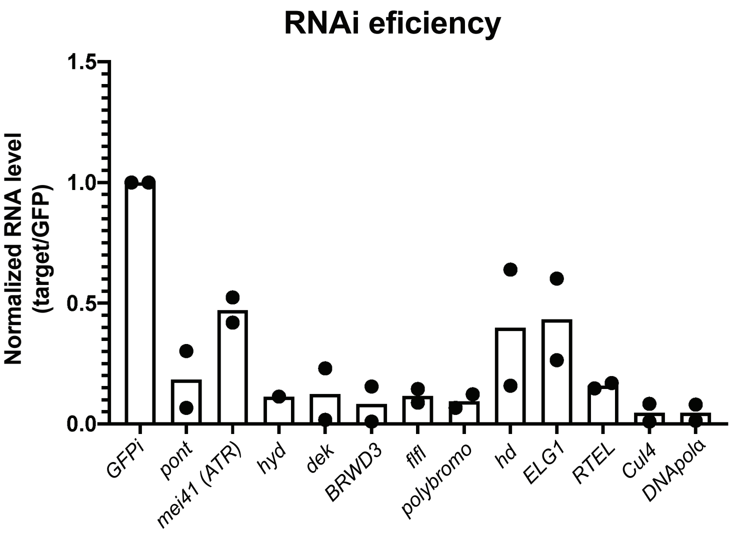


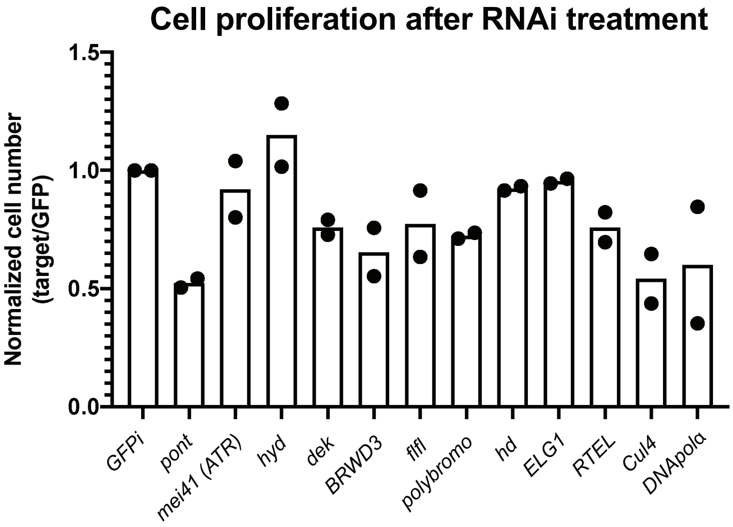


**Supplemental Figure 2:** Validation of RNAi-based depletion of targets. **(A)** Normalized depletion efficiency for two biological replicates. The normalized ratio is the target/*Tubulin* in the non-targeting *GFP* control divided by target/*Tubulin* in the RNAi-treated cells **(B)** Cell proliferation after five days of RNAi depletion relative to the *GFP* non-targeting control.
